# Diversity and evolution of amphibian pupil shapes

**DOI:** 10.1101/2021.08.15.456426

**Authors:** Kate N. Thomas, Caitlyn Rich, Rachel Quock, Jeffrey W. Streicher, David J. Gower, Ryan K. Schott, Matthew K. Fujita, Rayna C. Bell

## Abstract

Pupil constriction has important functional consequences for animal vision, yet the evolutionary mechanisms underlying diverse pupil sizes and shapes, often among animals that occupy optically similar environments, are poorly understood. We aimed to quantify the diversity and evolution of pupil shapes among amphibians and test for potential correlations to ecology based on functional hypotheses. Using photographs, we surveyed pupil shape and the orientation of the constricted pupil across adults of 1293 amphibian species, 72 families, and 3 orders, and additionally for larval life stages for all families of frogs and salamanders with a biphasic ontogeny. Pupil shape is exceptionally diverse in amphibians with evolutionary transitions throughout the amphibian tree of life. For amphibians with a biphasic life history, we found that pupils change in many species that occupy distinct habitats before and after metamorphosis. Finally, we found that non-elongated (round or diamond) constricted pupils were correlated with species inhabiting consistently dim light environments (burrowing and aquatic species) and that elongated pupils (vertical and horizontal) were more common in species with larger absolute eye sizes. We propose that amphibians provide a valuable group within which to explore the anatomical, physiological, optical, and ecological mechanisms underlying the evolution of pupil shape.

## Introduction

The ability to detect light and form images is an important sensory modality for most animals. Almost all animal phyla have evolved light-sensitive organs, ranging from eye-spots that simply detect the presence or absence of light to compound eyes that detect fast movement, providing a wide field of view and allowing images to be formed (Halder *et al*., 1995). Vertebrates and cephalopods have camera-type eyes, in which the aperture of the iris (the pupil) modulates the amount of light that enters the eye. In most species, iris muscles contract or relax to dilate or constrict the pupil in response to ambient light levels. This action dynamically changes the size of the aperture, enabling the individual to adjust the sensitivity and resolution of their visual system to current conditions, termed the pupillary light response (e.g., Douglas, 2018). The configuration of iris musculature determines the extent of constriction and the shape of the constricted pupil, ranging from the round and fixed pupils of most teleost fishes to the highly dynamic and complex pupil shapes of cephalopods (Douglas, 2018). The shape of the constricted pupil, in combination with other optical properties of the eye, will also reduce spherical aberration and can determine which wavelengths of light are focused on the retina (Kröger *et al*., 1999; Malmström & Kröger, 2006; Roth *et al*., 2009). Despite the clear functional consequences of pupil constriction for animal vision, the evolutionary mechanisms underlying diverse pupil sizes and shapes, often among animals that occupy optically similar environments, are poorly understood.

Pupil diversity in vertebrates includes non-elongated shapes (e.g., circular) and vertically or horizontally elongated shapes (Douglas, 2018). Some vertebrate groups exhibit little variation in constricted pupil shape: birds, turtles, and teleost fishes all have predominantly non-elongated, round constricted pupils, and all crocodilian pupils constrict to a vertical slit (reviewed in Douglas, 2018). By contrast, within mammals, squamates (lizards and snakes), and amphibians (frogs, salamanders, and caecilians) constricted pupils include all three orientations (Douglas, 2018), which may reflect the greater diversity of light environments these lineages occupy and their corresponding visual ecologies. For instance, vertical pupil constriction in elapid snakes (cobras, mambas and marine snakes) is correlated with diel activity and foraging mode: the constricted pupils of nocturnal species that are ambush predators are vertical, whereas those of diurnal species that are active foragers are circular (Brishoux *et al*., 2010). In mammals, pupil constriction is also correlated with activity period, with horizontally elongated and non-elongated pupils occurring in diurnal species and vertically elongated or slit pupils in nocturnal and crepuscular species (Mann, 1931). Elongated, slit-like pupil constrictions are also hypothesized to enhance vision in particular orientations but with conflicting evidence. For instance, vertically elongated pupils have been proposed to increase depth of field in a horizontal plane (e.g., Brishoux *et al*., 2010) or alternatively in a vertical plane (e.g., Hart *et al*., 2006; Banks *et al*., 2015). These hypotheses, however, have been explored in only a relatively small subset of the phylogenetic and ecological diversity of vertebrates. Here, we aim to quantify the diversity and evolution of pupil shapes among amphibians and test for potential correlations to ecology based on functional hypotheses.

Amphibians are a speciose (>8,300 extant species: AmphibiaWeb, 2021), diverse, and ecologically rich radiation with repeated evolutionary transitions in activity period and habitat that influence the light environments in which they are active, and have evolved, in. Although pupil shape has been studied in the context of species identification and systematics in some groups (e.g., Drewes, 1984; Glaw & Vences, 1997; Nuin & do Val, 2005; Rödel *et al*., 2009; Menzies & Riyanto, 2015), the evolutionary lability and functional consequences of different pupil shapes in amphibians are poorly understood. The limbless caecilian amphibians (order Gymnophiona; >200 extant species) are predominantly fossorial with greatly reduced visual systems, including eyes covered by skin and/or bone in many lineages (Mohun *et al*., 2010; Walls, 1942; Wake, 1985; Wilkinson, 1997). In even the most extensively developed eye of extant caecilians, the iris musculature is rudimentary (Mohun & Wilkinson, 2015) or absent (Himstedt, 1995), making changes in pupil size and shape unlikely (Douglas, 2018); consequently, in this study we mostly focus on frogs (order Anura, >7300 extant species) and salamanders (order Caudata, >700 extant species). A recent study characterized variation in absolute and relative eye size across all anuran families, and determined that frogs generally have large eyes relative to other vertebrates and that variation in adult eye size is associated with differences in habitat, activity period, and breeding ecology (Thomas *et al*., 2020). Variation in salamander eye size has not yet been quantified, but this lineage is also ecologically diverse with fully aquatic, arboreal, and fossorial species that likely differ substantially in visual ecology. Both frogs and salamanders are considered visual predators, and behavioral studies in both groups indicate that visual signals and coloration can play an important role in intraspecific communication (Jarger & Forester, 1993; Haddad & Giaretta, 1999; Hödel & Amezquita, 2001; Starnberger *et al*., 2014; Yovanovich *et al*., 2017). Likewise, both groups include species that are primarily diurnal, primarily nocturnal, or that are active under a range of light conditions (Anderson & Wiens, 2017). Consequently, both visual acuity and color discrimination may be important for many amphibian species in bright and/or dim light conditions (e.g., Toledo *et al*., 2007; Robertson & Greene, 2017). Furthermore, species that are active in both bright and dim light, and/or that have particularly large eyes, may rely on a large pupillary range to optimise visual performance relative to their surroundings because slit pupils allow a larger range of contraction (Walls, 1942).

Many amphibians have a biphasic ontogeny with an aquatic larval stage (termed tadpoles in frogs) and terrestrial adult life stages (e.g., McDiarmid & Altig, 1999), whereas others retain aquatic lifestyles as adults, have semi-terrestrial larvae, or develop without a larval life stage (termed direct development). During amphibian metamorphosis, dramatic morphological and physiological changes occur, including alterations to the visual system (Hoskins, 1990). Changes in eye-body scaling (Shrimpton *et al*., 2021) and lens shape (Sivak & Warburg, 1980; Sivak & Warburg, 1983) across ontogeny in frogs suggest that several structural aspects of the visual system adapt to both tadpole and adult visual requirements. Likewise, whole-eye differential expression of aquatic tadpoles versus terrestrial juvenile frogs (Schott *et al*., *in review*) demonstrates changes in a suite of visual genes related to eye and retinal development, light detection, lens crystallins, and phototransduction, indicating substantial decoupling between life stages at the level of gene expression. The biphasic ontogeny and shift between aquatic larval and terrestrial adult habitats in many amphibians is unique among tetrapods and thus presents the opportunity to investigate whether pupil shape is adaptively decoupled between life stages.

Here we survey and classify constricted pupil shape and orientation across adults of 1293 amphibian species, 72 families, and 3 orders, and additionally for larval life stages for all families of frogs and salamanders with a biphasic ontogeny. We first test the hypothesis that pupil shape changes across biphasic ontogeny in species that occupy distinct habitats before and after metamorphosis. Second, we identify evolutionary lineages with extensive pupil shape variation and quantify transition rates in pupil shape across the phylogeny. Finally, we test whether pupil shape exhibits correlated evolution with traits relevant to amphibian visual ecology. Specifically, we test whether (1) non-elongated pupils are correlated with inhabiting consistently dim light (aquatic or fossorial habitats) environments, (2) elongated pupils are associated with nocturnal and crepuscular activity, (3) vertically elongated pupils are correlated with navigating complex vertical (arboreal) habitats, and (4) elongated pupils are more common in species with large absolute eye size.

## Methods

### Species sampling and pupil classification

To assess the diversity of pupil shapes across amphibians, we searched online photo databases (primarily CalPhotos, https://calphotos.berkeley.edu) for images in which the eye and partially or fully constricted pupil shape was visible. Dilated pupils are circular, therefore we assumed pupils that were elongated in photos were at least partially constricted. For non-elongated pupils it is more challenging to determine whether the pupil is constricted from a photo but we relied on pupil size relative to eye size. We aimed to sample at least one species per family or sub-family of all currently recognized amphibian orders with externally visible eyes (72 families; Frost, 2021). When suitable images for a target family or species were not available on CalPhotos, we searched for photographs on other user-upload sites (e.g., Flickr), field guides, and primary literature. In a few instances, we relied on our personal photographs and field notes. Constricted pupil shapes (round, diamond, almond/oval, slit, triangle, tear) and orientations (non-elongated, horizontally elongated, vertically elongated) for each species were independently classified and reviewed by at least two observers (examples of shapes and axes of constrictions are in Figure 1, S1). Any discrepancies were flagged and resolved with the input of additional observers and photographs when available, or removed from the dataset when uncertainty remained. Larval frogs and salamanders, and adult caecilians, apparently lack or have a very weak pupillary response (Douglas, 2018) and thus our scoring in these instances are likely of permanently (or near-permanently) fixed pupil shapes that do not clearly correspond with either the fully dilated or fully constricted pupils of adult frogs and salamanders. Likewise, we note that almond or oval shapes in both horizontally and vertically elongated pupils may further constrict to a narrow slit under brighter light conditions. Because we relied on photographs to classify pupil shapes rather than experimentally assessing pupillary response, our determination of “almond or oval” versus “slit” pupil shapes were limited by the available photographs. However, our approach provides a more comprehensive survey of pupil constriction diversity in amphibians than is currently feasible with experimental approaches.

**Figure 1:**
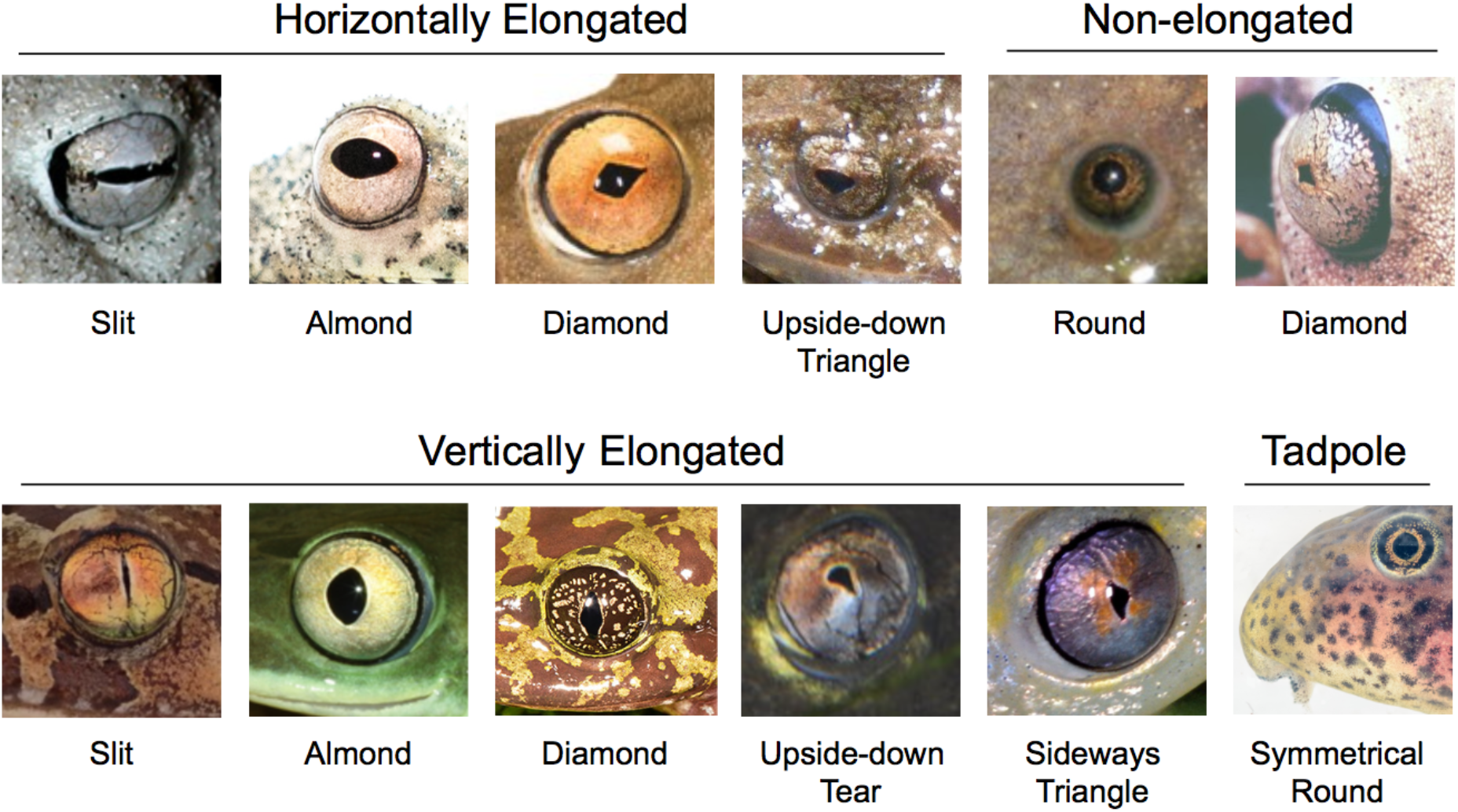
Examples of horizontally elongated, non-elongated, and vertically elongated adult and larval anuran pupil constriction shapes. We note that almond or oval pupil shapes may further constrict to a slit under bright light conditions. Photo credits (left to right, top to bottom) *Breviceps macrops* (Arie van der Meijden), *Hyperolius thomensis* (Andrew Stanbridge), *Boana boans* (Twan Lenders), and *Geocrinia lutea* (Grant Webster), *Xenopus tropicalis* (Daniel Portik), *Boana geographica* (Germano Woehl jr.), *Astylosternus batesi* (Greg Jongsma), *Tachycnemis seychellensis* (Gonçalo Rosa), *Heleophryne rosei* (Courtney Hundermark), *Calyptocephalella gayi* (Peter Janzen), and *Heterixalus betsileo* (Bernard Dupont), *Hylarana albolabris* (Christian Irian).

Once we had surveyed representatives of each family or sub-family, we expanded our sampling to encompass species that were the focus of recent studies on anuran visual biology (e.g., Thomas *et al*., 2020; Thomas *et al*., *in review*; Shrimpton *et al*., 2021) to maximize overlap with existing datasets. Preliminary assessments of this diversity suggested that pupil shape was diverse and/or evolutionarily labile in particular lineages and thus we elected to sample these groups in more depth. This included families in the Afrobatrachia radiation (Arthroleptidae, Brevicipitidae, Hemisotidae, Hyperoliidae), and the families or super-families Hylidae, Microhylidae, and Myobatrachidae. For families with extensive ecological diversity as adults (i.e., fully aquatic, semi-aquatic, ground-dwelling, arboreal, fossorial) we aimed to sample representative species of this diversity. Our final dataset included pupil shape observations for 1241 species of Anura (56 families), 43 species of Caudata (10 families), and 9 species of Gymnophiona (6 families). Pupil shape and associated references are given in the supplementary data files.

### Phylogeny

We used the phylogenetic hypothesis of Jetz & Pyron (2018) for visualizing trait distributions and modeling trait evolution across species. This phylogeny used a molecular backbone as well as taxonomic information to include proposed relationships among 7238 amphibian species. We matched the phylogeny to our dataset and performed all subsequent analyses using R v.4.1.0 (R Core Team, 2021) in RStudio v.1.4.1717 (RStudio Team, 2021). We used the R package AmphiNom v.1.0.1 (Liedtke, 2019) to match tip labels in the phylogeny to species names in our dataset by converting both to the taxonomy of Frost (2021) and manually checking and matching any species with multiple synonyms. For 46 species in our dataset not represented in the phylogeny, we used published literature to find the closest sister taxa that were represented in the tree (Table S1) and then added the missing species to the node representing the most recent common ancestor of these taxa using the getSisters, findMRCA, and bind.tip functions in phytools v.0.7.70 (Revell, 2012). Finally, we pruned the phylogeny to the 1293 species in our dataset using drop.tip in ape v.5.4.1 (Paradis *et al*., 2004; Paradis & Schliep, 2019) and randomly resolved polytomies with the bifurcr function in the PDcalc v.0.4.3.900 package (Nipperess & Wilson, 2021).

### Adult habitat and activity period classification

Adult ecology was categorized into binary states for activity pattern and different aspects of habitat using peer-reviewed literature, online natural-history resources, field guides, and field observations (see Supplemental References): 1) primarily diurnal or non-diurnal, 2) aquatic or non-aquatic, 3) fossorial or non-fossorial, and 4) scansorial or non-scansorial (see Supplemental data). Categorizations were simplified versions of those used by Thomas *et al*. (2020). Species were classed as primarily diurnal if adults are primarily active in daylight above ground; arrhythmic, cathemeral, crepuscular, and nocturnal species were all classified as non-diurnal. Species in which adults are primarily visually active underwater were categorized as aquatic. Species were classified as fossorial if adults are active underground, typically in soil (as opposed to only aestivating or sheltering underground). Finally, species in which adults climb up off the ground in vegetation were classified as scansorial.

### Pupil shape across biphasic ontogeny

To assess variation in pupil shape among larval frogs and salamanders, we searched through field guides, primary literature, and online photograph databases (e.g., CalPhotos, Flickr) and categorized pupils as described above. We classified larval pupil shape and orientation for at least one species in every family that has species with a larval life stage, including representative species with different larval habitats (i.e., semiterrestrial, phytotelm, pond, and stream-dwelling). To identify which lineages exhibit changes in pupil shape between larval and adult life stages, we classified pupil shape in adults for all species for which we determined larval pupil shape. As with larval habitat diversity, we also aimed to maximize adult habitat diversity in this paired sampling (i.e., fully aquatic, semi-aquatic, ground-dwelling, arboreal, fossorial). Both larval and adult habitat classifications were determined based on field guides, primary literature, and expert knowledge. To visualize variation in an evolutionary context, we mapped tadpole and adult pupil shapes and habitats on the modified Jetz & Pyron (2018) phylogeny using ape (Paradis *et al*., 2004; Paradis & Schliep, 2019). Pupil shape, habitat classifications, and associated references are listed in the supplementary data files.

### Evolutionary transitions of pupil orientation across the amphibian phylogeny

To gain insights into the evolutionary history and lability of adult pupil shape across the amphibian phylogeny, we implemented stochastic character mapping (Bollback, 2006) for the three categories of pupil orientation (non-elongated, vertically elongated, horizontally elongated). We used the fitDiscrete function in phytools v 0.7.70 (Revell, 2012) to fit equal-rates, symmetrical-rates, and all-rates-different models of character evolution. To select the “best” model, we compared AICc scores and AIC weights and then used make.simmap with the best-fit transition model (all-rates-different) to simulate character evolution across 100 trees. We plotted the phylogeny with branches colored based on the highest likelihood state of the node it originated from, and summarized mean pairwise transitions between each set of states across the 100 simulations.

### Effects of species ecology on pupil orientation

We implemented multivariate phylogenetic logistic regression in the R package phylolm (Paradis & Claude, 2002; Ives & Garland, 2010; Tung Ho & Ane, 2014) to examine the correlation structure among binary states for pupil orientation and ecology, using the general model format of pupil elongation ^~^ ecology. We tested three predictions in three separate models. First, we tested whether non-elongated pupils are associated with diurnal activity patterns (pupils: 0 = elongated, 1 = non-elongated; ecology: 0 = non-diurnal, 1 = diurnal). Second, we tested whether vertically elongated pupils are associated with scansoriality (pupils: 0 = non-vertical, 1 = vertical; ecology: 0 = non-scansorial, 1 = scansorial). Third, we tested whether non-elongated pupils are associated with fossorial and aquatic habitats (pupils: 0 = elongated, 1 = non-elongated; ecology: 0 = not fossorial or aquatic, 1 = fossorial or aquatic). We used the logistic_MPLE method, which maximizes the penalized likelihood of the logistic regression, and ran 1000 bootstrap replicates to estimate coefficients.

To test the prediction that species with large eyes would benefit from having a large pupillary range facilitated by elongated pupils, we tested for a correlation between eye size (transverse eye diameter) and elongated (horizontal or vertical) constricted pupils using a phylogenetic least squares regression in caper v.1.0.1 (Orme *et al*., 2018). Eye size data for 207 anuran species representing 54 families were gathered from a previous study (Thomas *et al*., 2020a; 2020b) and matched to our pupil dataset. We used phytools v.0.7.80 (Revell 2012) and ggplot2 v.3.3.3 (Wickham 2016) to visualize the data.

### Data availability statement

The datasets supporting this article are available from the Dryad Digital Repository: [provided upon acceptance, currently Supplemental Data], and code to replicate analyses and generate figures is available on GitHub ([provided upon acceptance]).

## Results

### Pupil shape diversity across amphibians

We examined pupil shape in nine species of Gymnophiona that occupy aquatic or fossorial habitats, all of which had non-elongated, circular pupils (Figure 2, S1). Pupil shape was more diverse across the 43 species of Caudata we classified, with non-elongated (circular) and horizontally elongated (almond/oval, slit, upside-down triangle) pupil shapes (Figure 2, S1). Our sampling of salamanders included a greater diversity of habitats than the caecilians (aquatic, fossorial, and scansorial) and also included diurnal species. Finally, we observed the greatest diversity of pupil shape in the 1241 species of Anura we examined, including non-elongated (circular, diamond), horizontally elongated (almond/oval, diamond, slit, upside-down triangle), and vertically elongated (almond/oval, diamond, sideways triangle, slit, upside-down teardrop) constricted pupils (Figure 1, 2). Our sampling of frogs included the greatest diversity of species and ecologies (i.e., fully aquatic, semi-aquatic, ground-dwelling, arboreal, fossorial) and we sampled representative species of this diversity within families. Pupil shape was notably diverse in the Afrobatrachia radiation (Arthroleptidae, Brevicipitidae, Hemisotidae, Hyperoliidae) and the families or super-families Hylidae, Microhylidae, and Myobatrachidae with all three major axes of pupil constriction represented in each of these lineages (Figure 2, S2). By contrast, other speciose and ecologically diverse lineages, such as Bufonidae, all exhibited horizontal almond/oval pupil shapes (Figure 2, S2). Adults of the fully-aquatic clawed frogs (Pipidae), giant salamanders (Cryptobranchidae), sirens (Sirenidae), amphiumas (Amphiumidae), and torrent salamanders (Rhyacotritonidae) all exhibited round pupils (Figure 2, S2) as did the fossorial frogs in Rhinophrynidae and Nasikabatrachidae (Figure 2, S2).

**Figure 2:**
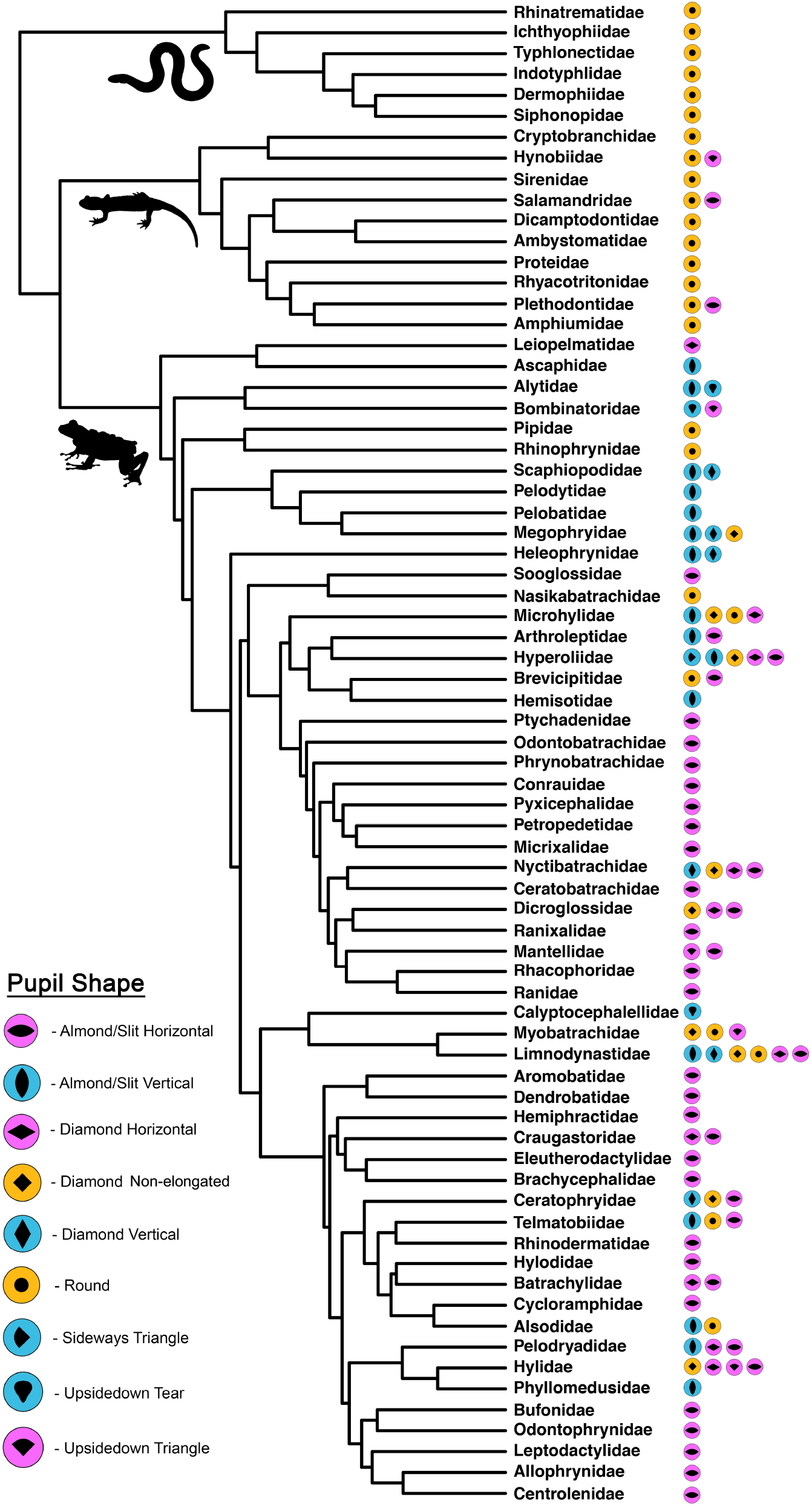
A phylogeny of all amphibian families that have developed eyes, showing the pupil shapes and constrictions we found within that family. We note that almond or oval pupil shapes may further constrict to a slit under bright light conditions but this was not always possible to assess from the available photographs. The phylogeny is modified from Jetz & Pyron (2018) and the complete dataset is shown in Figure S2.

### Pupil shape across biphasic ontogeny

Larval pupil shape was circular in all 92 species of frog and salamander that we surveyed regardless of their habitat (Figure 1, 3). For instance, the larvae of the Reed Frog *Hyperolius thomensis* develop in phytotelma (small pools of murky water that collect in tree cavities) and have non-elongated, circular pupils like those of the larvae of congeners *H. endjami*, which develop in ponds and streams (Figure S2, Supplementary data files). Likewise, semi-terrestrial tadpoles that develop in the splash zones of waterfalls (e.g., Rock River Frog *Thoropa miliaris*), in terrestrial nests (e.g., Nurse Frog *Allobates magnussoni*), or in dorsal pouches (e.g., Marsupial Frog *Gastrotheca piperata*) all have circular pupils. The only exception was the fossorial tadpoles of the Dancing Frog *Micrixalus herrei*, which hide within the gravel of streambeds, and appear to have skin-covered eyes as larvae but fully-developed eyes with horizontal pupils as adults (Senevirathne *et al*., 2016). In the 10 species (3 Anura and 7 Caudata) in our dataset that inhabit aquatic habitats as both larvae and adults, pupil shape remained circular in adults (Figure 3, S2). In the species that transition from an aquatic larval stage to a fossorial, scansorial, or ground-dwelling adult life stage, we observed all three axes of constricted pupil in adults (Figure 3, S2). Collectively, these observations indicate that pupil shape changes across biphasic ontogeny in many frog species that occupy distinct habitats before and after metamorphosis.

**Figure 3:**
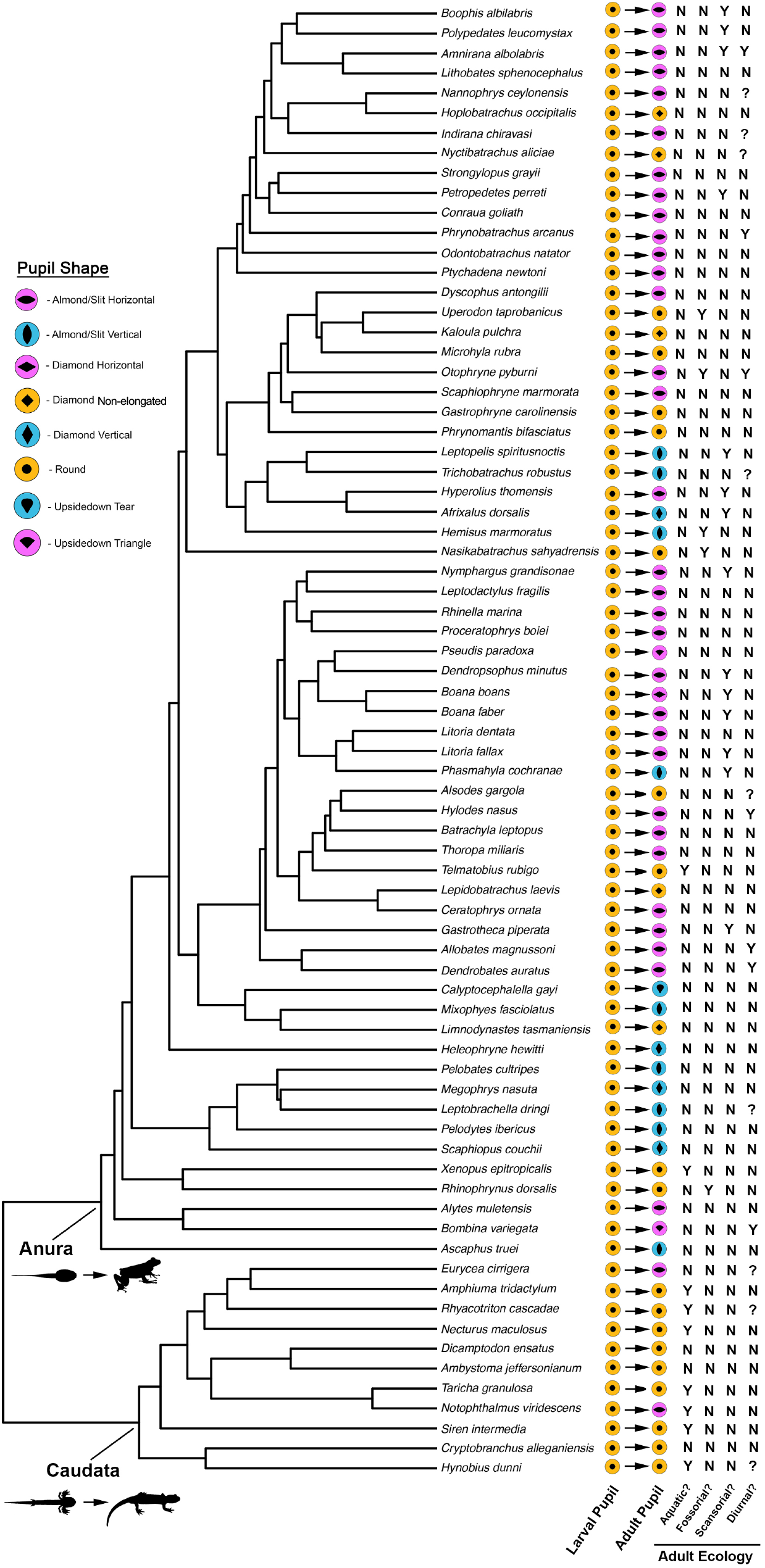
A phylogeny of larval and adult species pairs in our dataset (including representative species for all amphibian families that have a larval life stage with developed eyes) with observed pupil shapes and constrictions. The phylogeny is modified from Jetz & Pyron (2018) and the complete dataset is shown in Figure S2.

### Evolutionary transitions pupil orientation across the amphibian phylogeny

The “all-rates-different” model of character evolution was by far the best fit to our data for pupil orientation (Table 1) and we found high transition rates across the phylogeny (average of 84.56 changes between states across 100 total simulations) demonstrating the high evolutionary lability of this trait. The majority of transitions occurred from non-elongated to horizontally elongated pupils, whereas transitions from horizontally elongated to vertically elongated pupils were the least common (Figure 4). Many of the evolutionary transitions were concentrated within the Afrobatrachia radiation (Arthroleptidae, Brevicipitidae, Hemisotidae, Hyperoliidae) and the families or super-families Microhylidae and Myobatrachidae.

**Table 1:** Comparison of three Mk models of discrete character evolution for pupil constriction (non-elongated, vertically elongated, horizontally elongated) across sampled amphibian species (n = 1293). Models include an equal rates model with one transition rate parameter, a symmetric rates model with 3 transition rate parameters, and an all rates different model with 6 transition rate parameters.

| Model | log-lik | AICc | ΔAICc | AIC weight |
| --- | --- | --- | --- | --- |
| equal rates | -342.6 | 687.2 | 44.6 | 0 |
| symmetric rates | -339.7 | 685.4 | 41.9 | 0 |
| all rates different | -316.5 | 645.0 | 0 | 1 |

**Figure 4:**
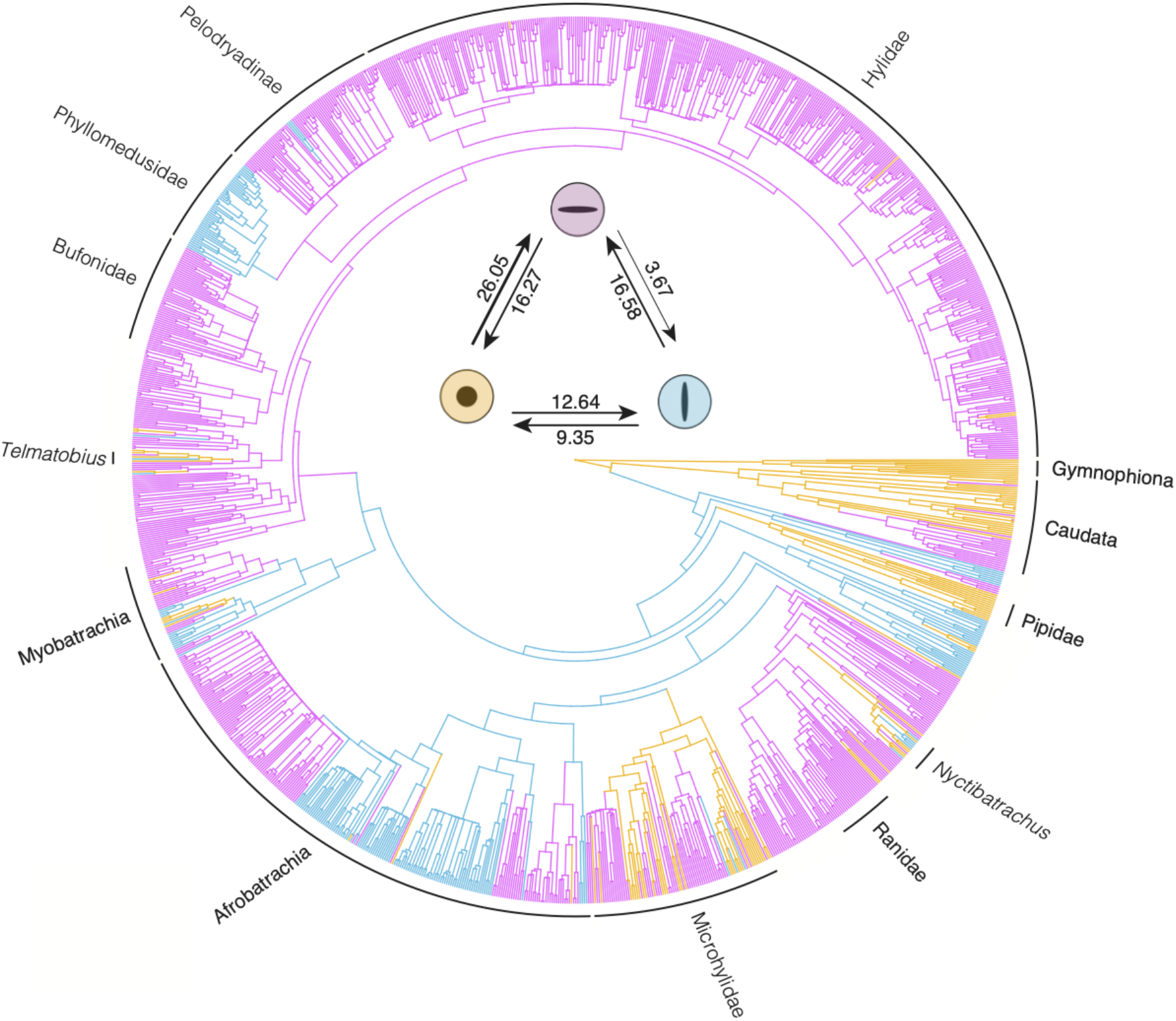
Distribution of non-elongated and elongated pupil constriction (or shape for Gymnophiona) in adult life stages of 1293 amphibian species (phylogeny modified from Jetz & Pyron 2018). Branches are colored by the highest probability state of the most recent node based on stochastic character mapping with an all-rates-different transition model across 100 trees. Lineages discussed in the text are labeled for reference. Inset depicts estimated transitions between non-elongated, horizontally elongated, and vertically elongated pupils based on stochastic character mapping. The thickness of the arrows is proportional to the mean transitions estimated across 100 simulations.

**Figure 5:**
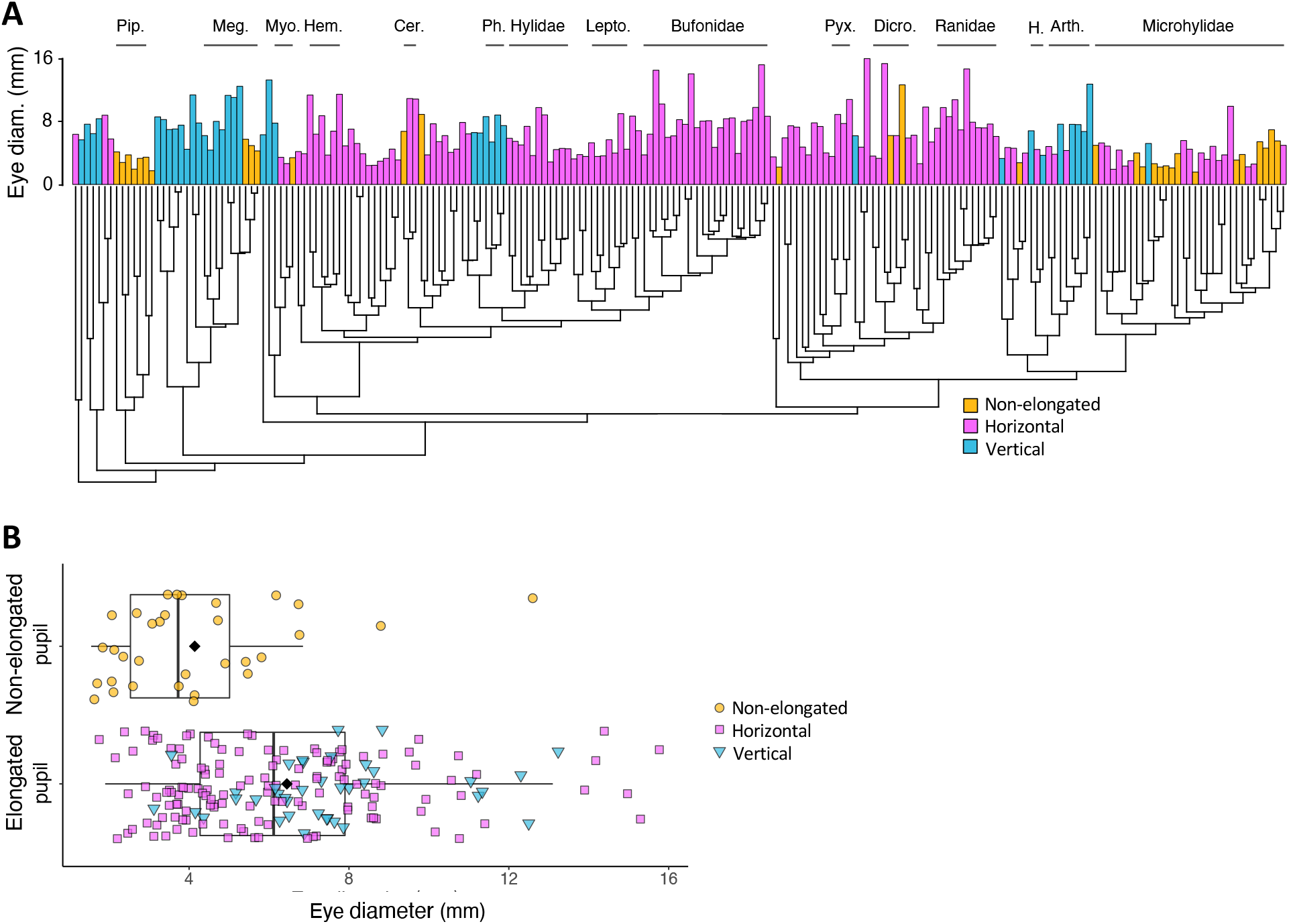
Eye size and the orientation of pupil constriction across 207 species of anuran amphibians (A). Species with elongated (horizontal or vertical) pupils have significantly larger eye diameters than those with non-elongated constricted pupils (B). Pip. = Pipidae, Meg. = Megophryidae, Myo. = Myobatrachidae, Hem. = Hemiphractidae, Cer. = Ceratophryidae, Ph. = Phyllomedusidae, Lepto. = Leptodactylidae, Pyx. = Pyxicephalidae, Dicro. = Dicroglossidae, H. = Hyperoliidae, Arth. = Arthroleptidae.

### Correlated evolution of species ecology and pupil orientation

Tests for correlation between the orientation of pupil constriction and ecology indicated that fossorial and aquatic ecologies are associated with non-elongated pupils in amphibians, while diurnality and scansoriality have no effects on pupil orientation (Table 2). Multivariate phylogenetic logistic regression found no significant association between non-elongated pupils and diurnal activity patterns across the 648 species in our activity pattern dataset; in fact, all of the 72 primarily diurnal species studied had horizontally or vertically elongated pupils. Likewise, we found no association between vertically elongated pupils and scansorial behaviors across 904 species. However, fossorial or aquatic habitats were a significant predictor of non-elongated pupils among 902 species tested (Table 1). Finally, among 207 anuran amphibians with data for both eye size and pupil shape, species with vertically or horizontally elongated pupils had significantly larger eyes than species with non-elongated pupils (PGLS: F = 5.24, df = 1 and 205, R^2^_adj_·= 0.02, p = 0.02).

**Table 2:** Summary of multivariate phylogenetic logistic regression analyses for the effects of binary ecological traits on binary pupil constriction orientation. Bolded predictors of pupil constriction are significant at Wald-type P < 0.05 for the given alpha value. Alpha is the phylogenetic correlation parameter estimate from phyloglm. Coefficient estimates are shown with upper and lower bootstrap estimates in parentheses based on 1000 fitted replicates.

| Model | $N_{\text{species}}$ | Alpha | Intercept | Predictor Coefficient |
| --- | --- | --- | --- | --- |
| Non-elongated pupil ~ |  |  | -1.52 | -0.07 |
| Diurnal activity | 648 | 0.004 | (-2.57, -0.27) | (-0.26, -0.62) |
| Vertical pupil ~ |  |  | -2.40 | 0.004 |
| Scansorial habitat | 904 | 0.005 | (-3.15, -0.30) | (-0.17, 0.14) |
| Non-elongated pupil ~ |  |  | **-2.06** | **2.23** |
| Aquatic/fossorial habitat | 902 | 0.005 | **(-3.39, -1.25)** | **(1.50, 3.39)** |

## Discussion

### Amphibians exhibit exceptional diversity of pupil shapes

In our assessment of pupil shape in nearly 1300 extant amphibian species (ca. 15% of described species) we observed great diversity among anurans including multiple shapes (e.g., circle, diamond, triangle, slit, teardrop) within each of the major axes of constriction (non-elongated, horizontally elongated, and vertically elongated). This diversity is in stark contrast to birds, turtles, and teleost fishes, which all have predominantly non-elongated, round constricted pupils and to crocodilians, in which pupils all constrict to a vertical slit (Douglas, 2018). Mammals and squamate reptiles, however, exhibit a wide diversity of pupil shapes, including shapes we have not observed in amphibians. For instance, many ungulates have horizontally constricting, rectangular pupil shapes (Miller & Murphy, 2016) that we did not observe in amphibians. Likewise, some geckos have scalloped edges along the pupil margin such that when the pupil constricts to a slit they are left with a vertical row of pinhole pupils (Mann, 1931). While we did not observe this extensive scalloping in amphibians, we did see irregular pupil margins in many anuran species (in association with opercula, elygia and umbracula e.g., brevicipetid Rain Frogs, centrolenid Glass Frogs) that could result in multiple pupil apertures if the pupil is further constricted than what we observed in available photographs. The proposed functional advantage of multiple apertures is that they enable accurate depth perception even when the pupil is constricted (Douglas, 2018). Alternatively, irregular pupil shapes may serve to conceal the eye as proposed for some bottom-dwelling fishes and for some reptiles (Walls, 1942; Douglas *et al*., 2002; Roth *et al*., 2009; Douglas, 2018; Youn *et al*., 2019). Finally, it has been suggested that pupil constriction matches the shape and location of increased photoreceptor density in the retina (i.e., retinal streaks) but this hypothesis is not supported in the birds, mammals, and fishes examined to date (Douglas, 2018). The variation in amphibian pupil shape we documented in the present study, particularly in frogs, warrants further attention with respect to the underlying musculature of the iris, latency and extent of the pupillary response, properties of the lens, and the arrangement of photoreceptor cells in the retina to better understand the functional consequences of this diversity.

### Ontogenetic changes in pupil shape

Our sampling of larval and adult pupil shape and constricted pupil orientation across 92 ecologically diverse species of frog and salamander indicates that pupils are likely non-elongated and round in most or all amphibian larvae. In addition, in many species that occupy distinct habitats before and after metamorphosis pupil shape changes during ontogeny. In particular, species that remain in aquatic habitats as adults retain non-elongated, round pupils whereas many species that occupy non-aquatic habitats as adults exhibit all three major axes of pupil constriction. Thus, our results are consistent with other studies of the visual system in larval and adult amphibians demonstrating that eye-body scaling (Shrimpton *et al*., 2021), lens shape (Sivak & Warburg, 1980; Sivak & Warburg, 1983), and whole-eye gene expression (Schott *et al*., *in review*) are decoupled when larvae and adults inhabit different light environments. Detailed examination of the iris musculature in developmental series of species that do and do not exhibit changes in pupil shape across ontogeny would provide greater insight into the key anatomical differences and onset of these changes within and among species. In addition, it is not clear whether pupils in some or all amphibian larvae have a pupillary light response. We explored this in larvae of two species (*Bufo bufo* and *Rana temporaria*) and did not observe any changes in pupil diameter or shape when exposed to bright light after 1 hour of dark adaptation (KNT, JWS, pers. obs.). We propose that future studies investigate the extent of pupillary response in a more diverse sample of amphibian larvae, including species that may experience a wider range of light environments than fully aquatic larvae (e.g., semi-terrestrial larvae).

### Transitions in pupil orientations across the phylogeny

Pupil shape is often considered an important diagnostic character for anuran systematics (e.g., Drewes, 1984; Nuin & do Val, 2005; Rödel *et al*., 2009; Menzies & Riyanto, 2015), and the orientation of pupil shape (non-elongated, horizontally elongated, vertically elongated) is largely conserved within several families that we extensively sampled (e.g., Bufonidae, Hylidae, Phyllomedusidae, Ranidae). Furthermore, pupil shape is conserved within (and divergent between) genera in some families (e.g., *Afrixalus* and *Hyperolius* in the family Hyperoliidae). Yet, we also found that some genera exhibited diversity in pupil orientation and shape among closely related species (e.g., *Nyctibatrachus*, *Telmatobius*). Thus, pupil shape appears to be an evolutionarily labile trait at both deep and recent timescales across the amphibia suggesting that this trait may not be a reliable character for systematics at some taxonomic levels in some lineages. Pupil shape also varies among closely related species in elapid snakes (Brischoux *et al*., 2010), and in felids and canids (Banks *et al*., 2015), likely reflecting the diverse visual environments these tetrapod groups occupy.

Ancestral character state reconstructions infer that the ancestral state for caecilians and salamanders was a non-elongated pupil whereas for frogs, vertically elongated pupils were the ancestral state. This result is consistent with the observation that teleost fishes exhibit predominantly non-elongated, circular constricted pupils (Douglas, 2018), and suggests that horizontal pupils evolved independently, and repeatedly, within salamanders and frogs. Elongated pupils are associated to some extent with multifocal lenses in which the lens has concentric zones of different focal lengths that enable the animal to correct for chromatic aberration or distortion (Malmström & Kröger, 2006). Consequently, an elongated pupil shape, which utilizes the whole lens diameter, enables the animal to use the full refractive range of the lens while regulating the total amount of light that enters the eye, thus providing crisp color vision in dim light (when the pupil is dilated and circular) and in bright light (when the pupil is constricted and elongated). The presence of elongated pupils in several anuran lineages, and in plethodontid and salamandrid salamanders, suggests they may have multifocal lenses and rely on color vision in a range of light environments (Malmström & Kröger, 2006), though multifocal lenses are also present in birds, which have circular pupils (Lind *et al*., 2008). Radiations like Afrobatrachia, which exhibit multiple transitions in the orientation of pupil elongation, diurnal and nocturnal activity periods, and include colorful and sexually dichromatic species (Portik *et al*., 2019), may be particularly fruitful for investigating the optical and evolutionary consequences of pupil elongation.

### Ecological correlates of pupil orientation in amphibians and other vertebrates

Animals that are active in a wide range of light levels, either because they are active both at nighttime and during the day or because they move between aquatic and terrestrial environments, tend to have a large pupillary range (Douglas, 2018). Pupils that are elongated (either vertically or horizontally) when constricted have a greater dynamic range than pupils that maintain a circular shape when constricted and are advantageous for species that rely on vision under a range of light conditions (Hart *et al*., 2006). Correspondingly, there was a significant correlation between non-elongated pupil constriction and amphibian species inhabiting consistently dimmer light environments (fully aquatic and fossorial lifestyles). Likewise, species with smaller absolute eye sizes tend to have non-elongated constricted pupils whereas those with larger absolute eye sizes have elongated pupils. However, there was no significant association between pupil elongation and activity period as proposed in other vertebrates such as snakes (Brishoux *et al*., 2010) and mammals (Mann, 1931). The vast majority of species in our dataset had elongated pupils, regardless of activity period and thus maintaining a greater range of pupil constriction is likely advantageous across most amphibian species.

Vertically elongated pupils are proposed to provide greater astigmatic depth of field in vertical planes (Banks, 2015), which could provide better spatial resolution for navigating complex vertical environments. However, we did not find a correlation between vertical pupils and scansorial behavior. Furthermore, horizontally elongated pupil constriction is prevalent across diverse families of largely arboreal species including hylid treefrogs and hyperoliid reed frogs. An alternative hypothesis for the functional consequences of vertically versus horizontally elongated pupils points to the visual ecology of predator versus prey species (Banks, 2015). In particular, vertical pupils are proposed to provide greater depth of field for ambush predators without the use of motion parallax movements whereas horizontally elongated pupils are proposed to improve image quality and provide greater field of view for detecting potential predators (Banks, 2015). Future studies of feeding ecology and predator avoidance in closely related species that differ in orientation of pupil shape may shed light on the functional consequences of vertically versus horizontally elongated pupils.

## Conclusion

Pupil shape is diverse in amphibians, especially in anurans, with evolutionary transitions throughout the amphibian tree of life. For amphibians with a biphasic life history, pupil shape changes in many species that occupy distinct habitats before and after metamorphosis, with all larvae having circular pupils. Furthermore, non-elongated pupils were correlated with species inhabiting consistently dim light environments (burrowing and aquatic species) and elongated pupils (vertical and horizontal) were more common in species with larger absolute eye sizes. We did not, however, find support for diurnal species having non-elongated pupil constriction or for species navigating complex vertical habitats (arboreal and scansorial) having vertically elongated pupils. Amphibians provide an exciting group for future research exploring the anatomical, physiological, optical, and ecological mechanisms underlying the evolution of pupil diversity.

## Acknowledgements

We thank R. Martins, I. Prates, G. Webster and the many amphibian researchers and enthusiasts who have shared their photographs in field guides and online databases. This study would not have been possible without their efforts. R. Douglas provided valuable guidance throughout the project and a thorough edit of the initial manuscript. H. Christoph Liedtke graciously shared code used in generating the figures. This work was supported by grants from the Natural Environment Research Council, UK (grant no. NE/R002150/1) and the National Science Foundation, USA (grant no. DEB #1655751).

